# Analysis of ISCB honorees and keynotes reveals disparities

**DOI:** 10.1101/2020.04.14.927251

**Authors:** Trang T. Le, Daniel S. Himmelstein, Ariel A. Hippen Anderson, Matthew R. Gazzara, Casey S. Greene

## Abstract

Delivering a keynote talk at a conference organized by a scientific society, or being named as a fellow by such a society, indicates that a scientist is held in high regard by their colleagues. To explore if the distribution of such indicators of esteem in the field of bioinformatics reflects the composition of this field, we compared the gender, name-origin, country of affiliation and race/ethnicity of 412 researchers who had been recognized by the *International Society for Computational Biology* (75 fellows and 337 keynote speakers) with over 170,000 researchers who had been the last authors on computational biology papers between 1993 and 2019. The proportion of female fellows and keynote speakers was similar to that of the field overall. However, names of East Asian origin have been persistently underrepresented among fellows and keynote speakers. Moreover, fellows and keynote speakers with an affiliation in the United States were overrepresented by a factor of 2.1; almost two thirds of this excess was accounted for by a deficit of 101 fellows and keynote authors from China, India, France and Italy. Within the US, we found an excess of white fellows and keynote speakers and a depletion of Asian fellows and keynote speakers.

## Introduction

Scientists’ roles in society include identifying important topics of study, undertaking an investigation of those topics, and disseminating their findings broadly. The scientific enterprise is largely self-governing: scientists act as peer reviewers on papers and grants, comprise hiring committees in academia, make tenure decisions, and select which applicants will be admitted to doctoral programs. A lack of diversity in science could lead to pernicious biases that hamper the extent to which scientific findings are relevant to minoritized communities. Furthermore, even though minoritized groups innovate at higher rates, their novel contributions are discounted [1]. One first step to address this systemic issue is to directly examine peer recognition in different scientific fields.

Gender bias among conference speakers has been recognized as an area that can be improved with targeted interventions [2,3,4,5]. Having more female organizers on conference committees is associated with having more female speakers [6]. At medical conferences in the US and Canada, the proportion of female speakers is increasing at a modest rate [7]. Gender bias appears to also influence funding decisions: an examination of scoring of proposals in Canada found that reviewers asked to assess the science produced a smaller gender gap in scoring than reviewers asked to assess the applicant [8].

Challenges extend beyond gender: an analysis of awards at the NIH found that proposals by Asian, black or African-American applicants were less likely to be funded than those by white applicants [9]. There are also potential interaction effects between gender and race or ethnicity that may particularly affect women of color’s efforts to gain NIH funding [10]. Another recent analysis found that minority scientists tend to apply for awards on topics with lower success rates [11]. This finding might be the result of minority scientists selecting topics in more poorly funded areas. Alternatively, reviewing scientists may not recognize the scientific importance of these topics, which may be of particular interest to minority scientists.

We sought to understand the extent to which honors and high-profile speaking invitations were distributed equitably among gender, race/ethnicity, and name origin groups by an international society and its associated meetings. As computational biologists, we focused on the <u>International Society for Computational Biology</u> (ISCB), its honorary Fellows as well as its affiliated international meetings that aim to have a global reach: <u>Intelligent Systems for Molecular Biology</u> (ISMB) and <u>Research in Computational Molecular Biology</u> (RECOMB).

We used multiple methods to predict the gender, race/ethnicity, and name origins of honorees. Existing methods were relatively US-centric because most of the data was derived in whole or in part from the US Census. We scraped more than 700,000 entries from English-language Wikipedia that contained nationality information to complement these existing methods and built multiple machine learning classifiers to predict name origin. We also examined the last authors for more than 170,000 computational biology publications to establish a field-specific baseline using the same metrics. The results were consistent across all approaches: we found a dearth of non-white speakers and honorees. The lack of Asian scientists among keynote speakers and Fellows was particularly pronounced when compared against the field-specific background.

## Results

We curated a dataset of ISCB honorees that included 412 honorees who were keynote speakers at international ISCB-associated conferences (ISMB and RECOMB) as well as ISCB Fellows. The ISCB Fellows set contained the complete set of Fellows named (2009–2019). Keynote speakers were available for ISMB for all years from 1993–2019. Keynote speakers for RECOMB were available for all years from 1997–2019. We included individuals who were honored multiple times as separate entries. For example, Christine Orengo was a keynote speaker at RECOMB 2004 and became an ISCB Fellow in 2016, and thus was counted twice in this list.

We sought to compare this dataset with a background distribution of potential speakers, which we considered to be last authors of bioinformatics and computational biology manuscripts. We scraped PubMed for manuscripts written in English from 1993–2019 with the MeSH term <u>“computational biology”</u>. We downloaded the metadata of manuscripts published in these journals from PubMed, which provided 173,735 articles for evaluation. For each article, we extracted its last author’s fore name and last name for analysis.

In a <u>previous version</u> of this work, instead of last authors, we examined corresponding authors of articles in three well-recognized computational biology journals, and the results were consistent with our findings here [12].

### Similar gender proportion between ISCB’s honorees and the field

We observed a gradual increase of the proportion of predicted female authors, arriving at 28.8% in 2019 (Fig. 1, left). In recent years, ISCB Fellows and keynote speakers appear to have similar gender proportion compared to the population of authors published in computational biology and bioinformatics journals (Fig. 1, right); however, it has not yet reached parity. Examining each honor category, we observed an increasing trend of honorees who were women, especially in the group of ISCB Fellows (see <u>notebook</u>), which markedly increased after 2015. Through 2019, there were a number of years when meetings or ISCB Fellow classes have a high probability of recognizing only male honorees and none that appeared to have exclusively female honorees. We sought to examine whether or not there was a difference in the proportion of female names between authors and honorees. A multiple linear regression of this proportion for the groups and year did not reveal a significant difference (*p* = 0.558).

**Figure 1:**
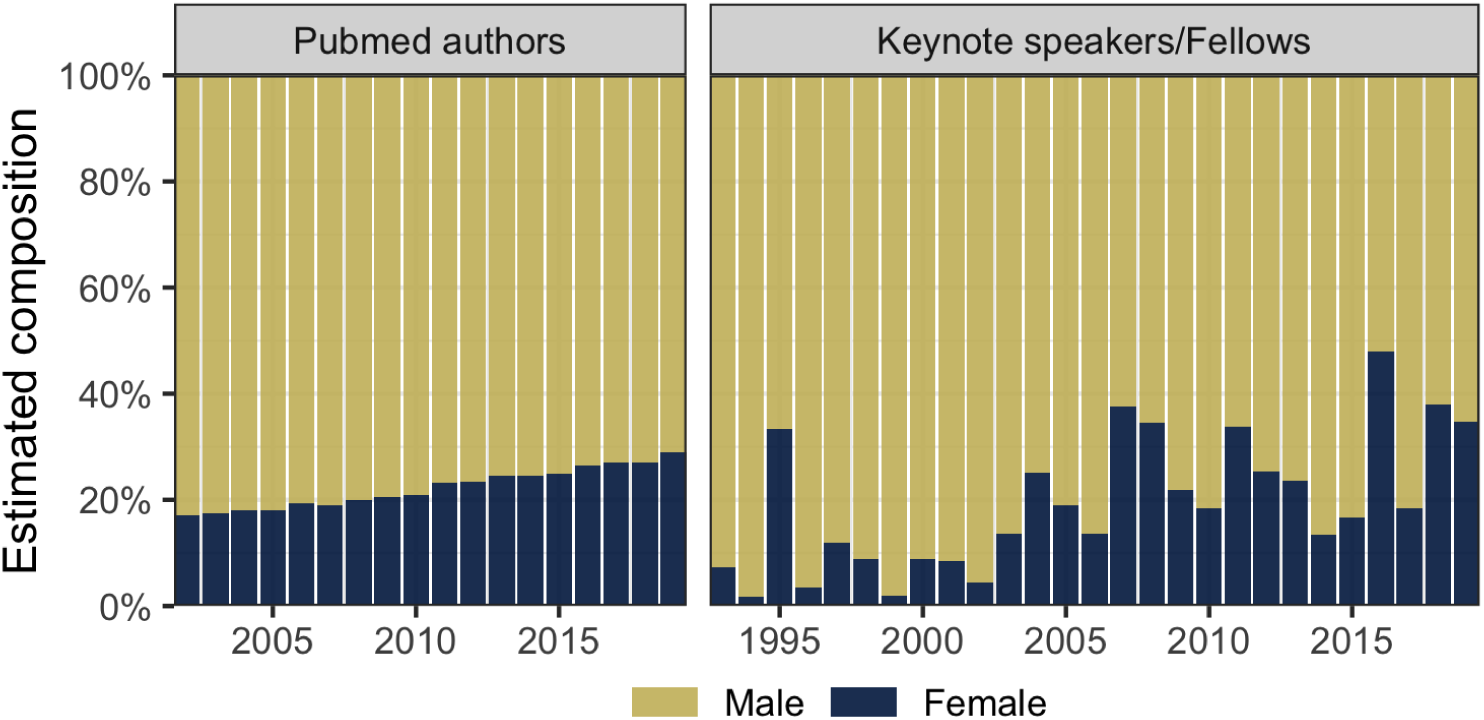
Estimated composition of gender prediction over the years of all Pubmed computational biology and bioinformatics journal authors (left), and all ISCB Fellows and keynote speakers (right). Male proportion (yellow) was computed as the average of the probability of being male of last authors (weight accordingly) or ISCB honorees each year. Female proportion (blue) was the complement of the male proportion. ISCB Fellows and keynote speakers appear have similar gender proportion compared to that of PubMed authors, but the proportion has not reached parity.

### Honorees with Celtic/English names are overrepresented while honorees with East Asian names are underrepresented

We applied our Wiki2019-LSTM model to both our computational biology honorees dataset and our dataset of last authors. We found that the proportion of authors with Celtic/English names had decreased (Fig. 2A, left). Among keynote speakers and fellows, we found that the majority of honorees are predicted to have Celtic/English or European names (Fig. 2A, right). When we directly compared honoree composition with PubMed, we observed discrepancies between the two groups (Fig. 2B). A name coming from the group of honorees has significantly higher probability of being Celtic/English (*β* _Celtic/English_ = 0.11924, *p* < 10^−9^) and lower probability of being East Asian (*β*_East Asian_ = −0.14791, *p* < 10^−9^). The two groups of scientists did not have a significant association with names predicted to be European and in Other categories (*p* = 0.39475 and *p* = 0.48625, respectively).

**Figure 2:**
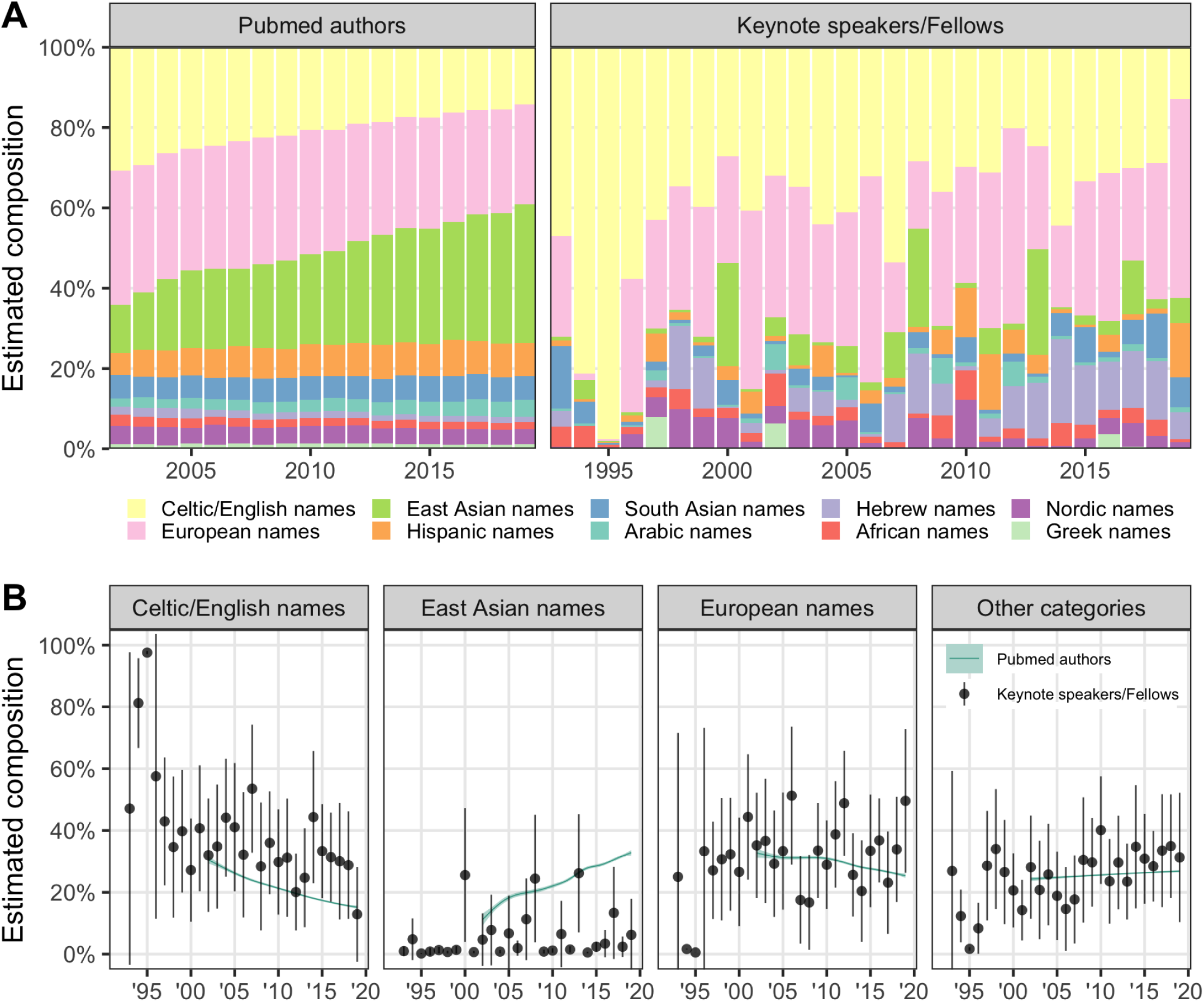
(A) Estimated composition of name origin prediction over the years of all Pubmed computational biology and bioinformatics journal authors (left), and all ISCB Fellows and keynote speakers (right). (B) For each region, the mean predicted probability of Pubmed articles is shown as teal GAM curve, and the mean probability and 95% confidence interval of the ISCB honoree predictions are shown as dark circles and vertical lines. Compared to the name collection of Pubmed authors, honorees with Celtic/English names are overrepresented while honorees with East Asian names are underrepresented. No statistically significant difference was observed between the proportion of honorees and authors with European names or in other categories (see Table 1).

We reached similar conclusion when applying the Wiki2019-LSTM model to the name origins of only US-affiliated scientists. We note that the US was not included in the training of the Wiki2019-LSTM model (see Methods). We found an overrepresentation of honorees with Celtic/English names (*β*_Celtic/English_ = 0.053, *p* = 0.031), a smaller overrepresentation of honorees with European names (*β*_European_ = 0.046, *p* = 0.042) and substantial underrepresentation of honorees with East Asian names (*β*_East Asian_ = −0.010, *p* = 3.6×10^−5^). No statistically significant difference was observed between the proportion of honorees and authors with names in Other categories (see Table 1, *p* = 0.95). Please see <u>analysis notebook</u> for more details.

**Table 1:**
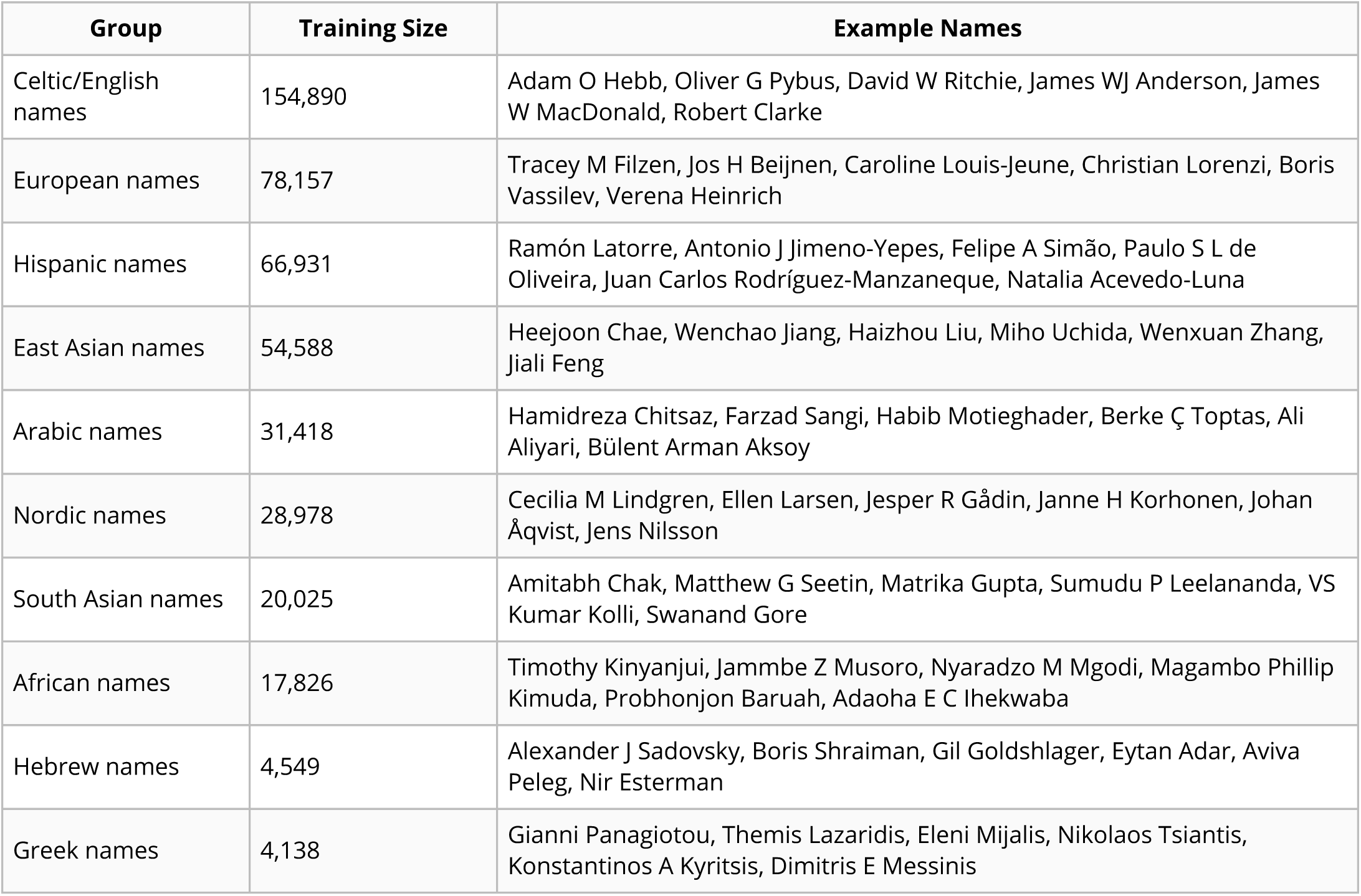
Predicting name-origin groups of names trained on Wikipedia’s living people. The table lists the 10 groups and the number of living people for each region that the LSTM was trained on. Example names shows actual author names that received a high prediction for each region. Full information about which countries comprised each region can be found in the online dataset <u>country_to_region.tsv</u>.

### Overrepresentation of US-affiliated honorees

We analyzed the countries of affiliation between last authors and ISCB honorees. For each country, we report a value of log_2_ enrichment (LOE) and its 95% confidence intervals. The full table with all countries and their corresponding enrichment can be browsed interactively in the corresponding <u>analysis notebook</u>. A positive value of LOE indicates a higher proportion of honorees affiliated with that country compared to authors. A LOE value of 1 represents a one-fold enrichment (i.e., observed number of honorees is twice as much as expected). In the 20 countries with the most publications, we found an overrepresentation of honorees affiliated with institutions and companies in the US (153 speakers more than expected, LOE = 1.1, 95% CI (0.9, 1.2)) and Israel (14 speakers more than expected, LOR = 2.5, 95% CI (1.7, 3.1)), and an underrepresentation of honorees affiliated with those in China, France, Italy, India, the Netherlands, Korea, Brazil and Taiwan (Fig. 3).

**Figure 3:**
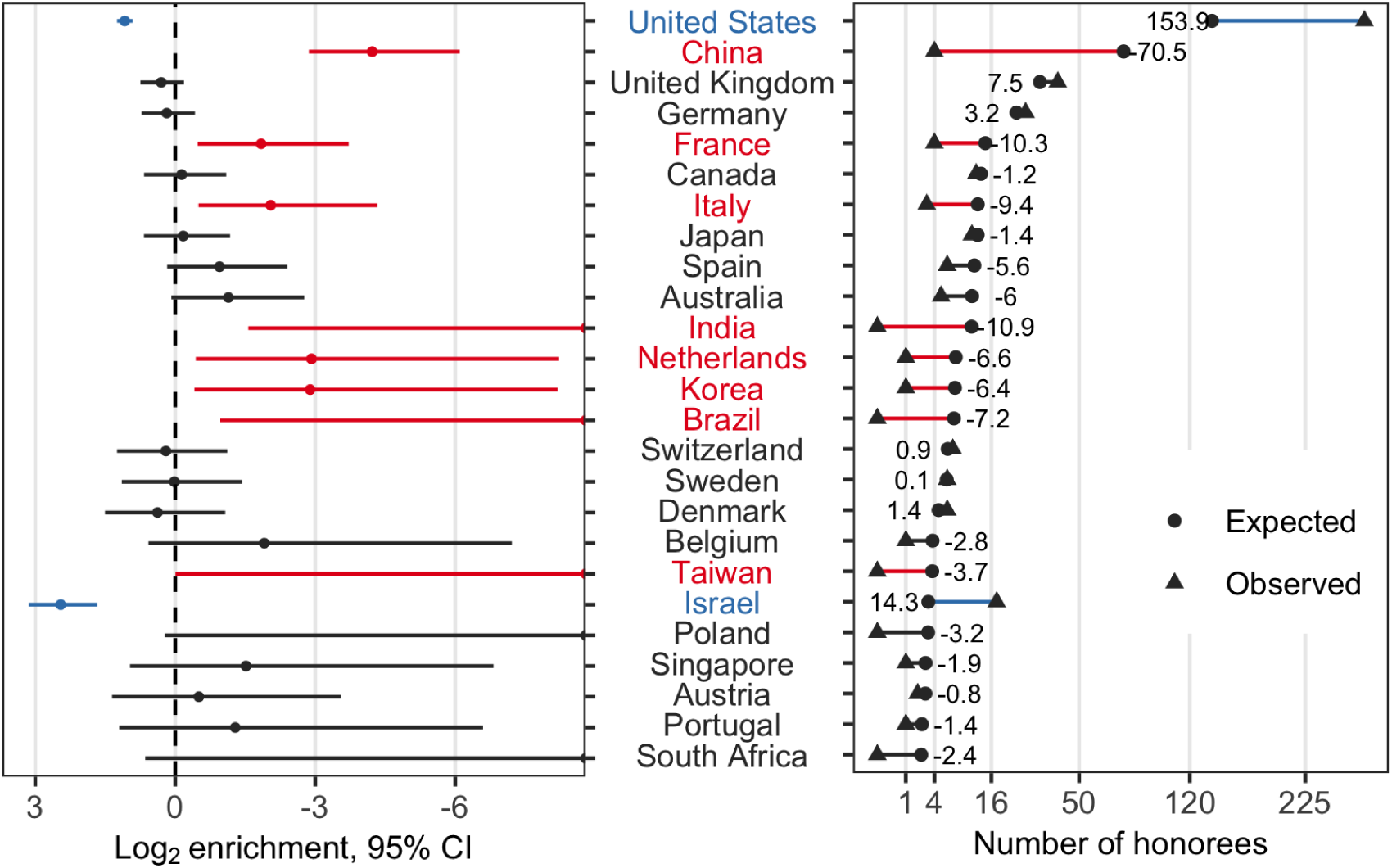
Each country’s log_2_ enrichment (LOE) and its 95% confidence interval (left), and the absolute difference between observed (triangle) and expected (circle) number of honorees (right). Positive value of LOE indicates a higher proportion of honorees affiliated with that country compared to authors. Countries are ordered based on the proportion of authors in the field. The overrepresentation of honorees affiliated with institutions and companies in the US and Israel contrasts the underrepresentation of honorees affiliated with those in China, France, Italy, India, the Netherlands, Korea, Brazil and Taiwan.

### Overrepresentation of white and underrepresentation of Asian honorees among US-affiliated scientists

We predicted the race and ethnicity of US-affiliated authors and honorees using wru, which is based on US census data. We found that an increasing proportion of last authors of computational biology papers whose last names associated with selecting Asian as a race/ethnicity category in the US census (Fig. 4A left). Compared to Pubmed authors, ISCB honorees with US affiliations have a higher proportion of individuals whose last names were associated with selecting white as a race/ethnicity category in the US census (Fig. 4A). Specifically, a name coming from the group of honorees has significantly higher probability of being white (*β*_white_ = 0.0975, *p* = 1.4×10^−5^) and lower probability of being Asian, (*β*_Asian_ = - 0.1122, *p* = 1.7×10^−6^). The two groups of scientists did not have a significant association with names predicted to be in Other categories (*p* = 0.16662). Separating honoree results by honor category did not reveal any clear differences among the categories (see <u>notebook</u>.

**Figure 4:**
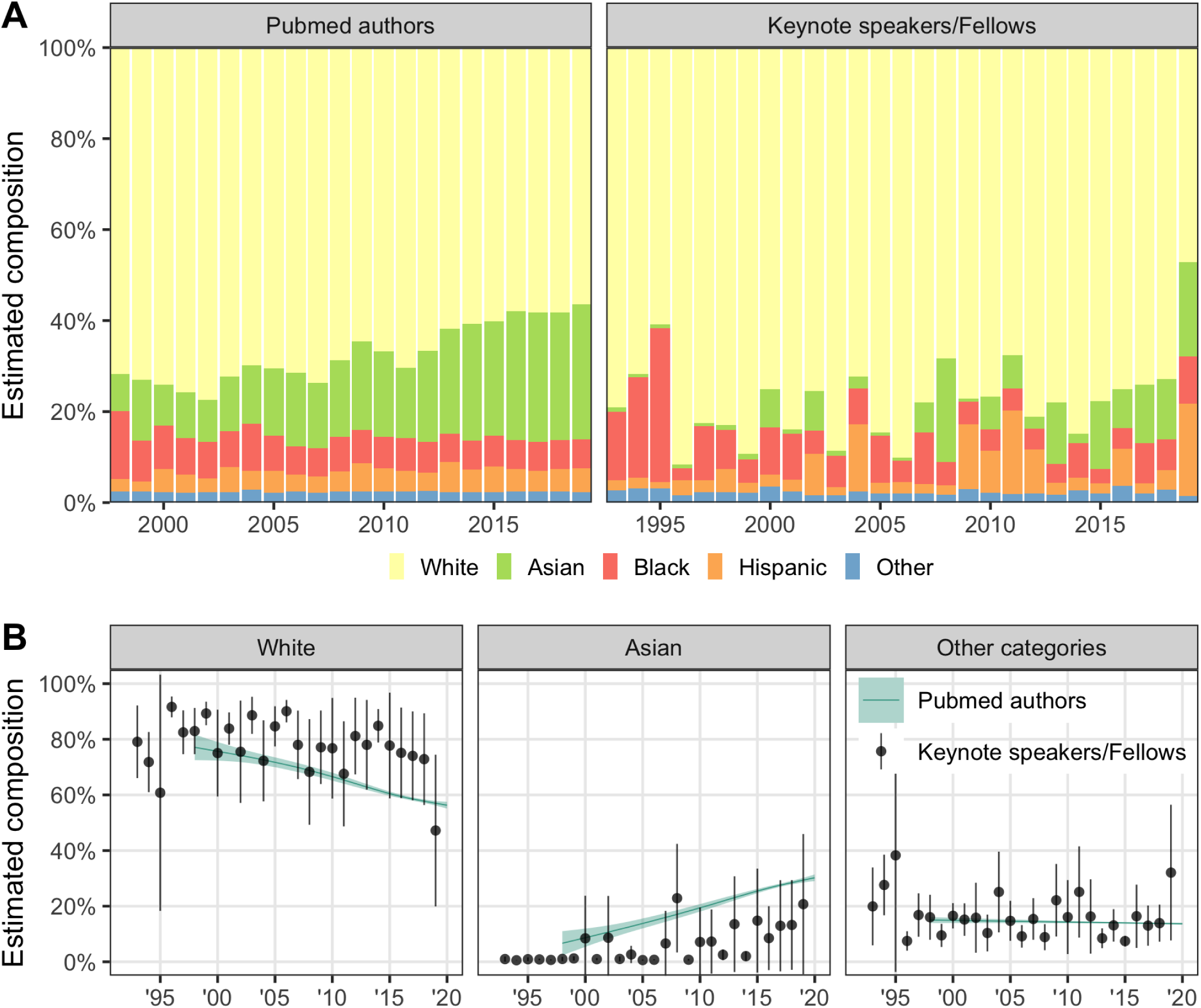
Estimated composition of census-based race/ethnicity prediction over the years of (A) all Pubmed computational biology and bioinformatics journal authors(left) and all ISCB Fellows and keynote speakers (right), (B) For each race/ethnicity category, the mean predicted probability of Pubmed articles is shown as teal generalized additive model (GAM) curve, and the mean probability and 95% confidence interval of the ISCB honoree predictions are shown as dark circles and vertical lines. The large difference between the means and the lack of overlaping of the confidence intervals in most years indicates an overrepresentation of white and underrepresentation of Asian honorees as compared to authors.

### Improvements to Honoree Diversity Subsequent to Our Primary Analysis

While our study was primarily designed to assess the diversity of honorees, the findings raise important questions about what can be done to address the disparities. By publishing our original findings on the biorxiv preprint server, we could begin to answer these questions by examining changes subsequent to our initial report. We released <u>version 1.0</u> of our manuscript on 2020-01-30. Early indications suggest ISCB is now increasing the diversity of honorees. In 2020, among 12 ISCB Fellows and 5 ISMB keynote speakers, the mean predicted probability of each honoree having an East Asian name is 33%, higher than any estimate in previous years (see <u>notebook</u>). The set of honorees also includes the first ISCB Fellow from China. Compared to past years, the 2020 honorees appear to better reflect the diversity of scientists in the computational biology field. These new results suggest: 1) deserving honorees who were members of under-recognized groups existed but had not been recognized until now, and 2) simply examining honoree distribution’s alignment with the field can trigger changes that address issues of unequal representation. However, we note that this analysis dealt only with more senior scientists (the last authors on scientific manuscripts) in the context of honors and that many years of changed honoree distributions will be required for the set of honored scientists to better reflect the field’s senior author contributions.

## Discussion

A major challenge that we faced in carrying out this work was to narrow down geographic origins for some groups of names. Some name origin groups, such as Hispanic names, are geographically disparate. We were unable to construct a classifier that could distinguish between names from Iberian countries (Spain and Portugal) from those in Latin America in the group of Hispanic names. Discrepancies in representation between these groups are thus undetectable by our classifier. Honoree counts of those with Hispanic names are influenced from Spain as well as Latin America. In such cases, our analyses may substantially understate the extent to which minoritized scientists are underrepresented among honorees and authors.

Biases in authorship practices may also result in our underestimation of the composition of minoritized scientists within the field. We estimated the composition of the field using corresponding author status, but in neuroscience [13] and other disciplines [14] women are underrepresented among such authors. Such an effect would cause us to underestimate the number of women in the field. Though this effect has been studied with respect to gender, we are not aware of similar work examining race, ethnicity, or name origins.

We acknowledged that our supervised learning approaches are neither error free nor bias free. Because wru was trained on the US census to make predictions, many of the missing predictions are on names not observed in the US census. Although the underestimation of the proportion of these names could not be quantified in the list of authors and honorees, we complemented this race/ethnicity prediction method with an additional name origin analysis. By integrating different methods and preserving uncertainty by analyzing prediction probabilities rather than applying a hard assignment for each prediction, we hope to alleviate these biases and discover insightful findings that correctly reflect the current representation diversity at conferences.

Focusing on an international society and meetings, we measured honor and authorship rates worldwide. In this setting, we observe disparities by name groups. Because invitation and honor patterns could be driven by biases associated with name groups, geography, or other factors, we cross-referenced name group predictions with author affiliations to disentangle the relationship between geographic regions, name groups and invitation probabilities. We found that disparities persisted even within the group of honorees with a US affiliation.

An important questions to ask when measuring representation is what the right level of representation is. We suggest that considering equity may be more appropriate than strictly diversity. For example, we found similar representation of women between authors and honorees, which suggests honoree diversity is similar to that of authors. However, if fewer women are in the field because of systemic factors that inhibit their participation, we would not have reached equity. In addition to holding fewer corresponding authorship positions, on average, female scientists of different disciplines are cited less often [15], invited by journals to submit papers less often [14], suggested as reviewers less often [18], and receive significantly worse review scores [16]. Meanwhile, a review of women’s underrepresentation in math-intensive fields argued that today’s underrepresentation is not explained by historic forms of discrimination but factors surrounding fertility decisions and lifestyle choices, whether freely made or constrained by biology and society [19]. A recent analysis of gender inequality across different disciplines showed that, although both gender groups have equivalent annual productivity, women scientists have higher dropout rates throughout their scientific careers [20].

We found that ISCB’s honorees and keynote speakers, though not yet reaching gender parity, appear to have similar gender proportion to the field as a whole. On the other hand, honorees include significantly fewer people of color than the field as a whole, and Asian scientists are dramatically underrepresented among honorees. Societies’ honoree selection process failing to reflect the diversity of the field can play a part in why minoritized scientists’ innovations are discounted [1]. Although we estimate the fraction of non-white and non-Asian authors to be relatively similar to the estimated honoree rate, we note that both are represented at levels substantially lower than in the US population.

Societies, both through their honorees and the individuals who deliver keynotes at their meetings, can play a positive role in improving the presence of female STEM role models, which can boost young students’ interests in STEM [19] and, for example, lead to higher persistence for undergraduate women in geoscience [21]. Efforts are underway to create Wikipedia entries for more female [22] and black, Asian, and minority scientists [23], which can help early-career scientists identify role models. Societies can contribute toward equity if they design policies to honor scientists in ways that counter these biases such as ensuring diversity in the selection committees.

The central role that scientists play in evaluating each other and each other’s findings makes equity critical. Even many nominally objective methods of assessing excellence (e.g., h-index, grant funding obtained, number of high-impact peer-reviewed publications, and total number of peer-reviewed publications) are subject to the bias of peers during review. These could be affected by explicit biases, implicit biases, or pernicious biases in which a reviewer might consider a path of inquiry, as opposed to an individual, to be more or less meritorious based on the reviewer’s own background [11]. Our efforts to measure the diversity of honorees in an international society suggests that, while a focus on gender parity may be improving some aspects of diversity among honorees, contributions from scientists of color are underrecognized.

## Materials and Methods

### Honoree Curation

From <u>ISCB’s webpage listing **ISCB Distinguished Fellows**</u>, we found recipients listed by their full names for the years 2009–2019. We gleaned the full name of the Fellow as well as the year in which they received the honor. We identified major ISCB-associated conferences as those designated flagship (ISMB) or those that had been held on many continents (RECOMB). To identify **ISMB Keynote Speakers**, we examined the webpage for each ISMB meeting. The invited speakers at ISMB before 2001 were listed in the Preface pages of each year’s proceedings, which were archived in the <u>ISMB collection</u> of the AAAI digital library. We found full names of all keynote speakers for the years 1993–2019.

For the RECOMB meeting, we found conference webpages with keynote speakers for 1999, 2000, 2001, 2004, 2007, 2008, and 2010–2019. We were able to fill in the missing years using information from the RECOMB 2016 proceedings, which summarizes the first 20 years of the RECOMB conference [25]. This volume has two tables of keynote speakers from 1997–2006 (Table 14, page XXVII) and 2007–2016 (Table 4, page 8). Using these tables to verify the conference speaker lists, we arrived at two special instances of inclusion/exclusion. Although Jun Wang was not included in these tables, we were able to confirm that he was a keynote speaker in 2011 with the RECOMB 2011 proceedings [26], and thus we included this speaker in the dataset. Marian Walhout was invited as a keynote speaker but had to <u>cancel</u> the talk due to other obligations. Because her name was neither mentioned in the 2015 proceedings [27] nor in the above-mentioned tables, we excluded this speaker from our dataset.

### Name processing

When extracting honoree names, we began with the full name as provided on the site. Because our prediction methods required separated first and last names, we chose the first non-initial name as the first name and the final name as the last name. We did not consider a hyphen to be a name separator: for hyphenated names, all components were included. For metadata from PubMed and PMC where first (fore) and last names are coded separately, we applied the same cleaning steps. We created <u>functions to simplify names</u> in the pubmedpy Python package to support standardized fore and last name processing.

### Last author extraction

We assumed that, in the list of authors for a specific paper, last authors (often research advisors) would be most likely to be invited for keynotes or to be honored as Fellows. Therefore, we utilized <u>PubMed</u> to retrieve last author names to assess the composition of the field. PubMed is a search engine resource provided by the US National Library of Medicine and index scholarly articles. PubMed contains a record for every article published in journals it indexes (30 million records total circa 2020), within which we were able to extract author first and last names and their order using the E-Utilities APIs. To automate and generalize these tasks, we created the <u>pubmedpy</u> Python package.

From PubMed, we compiled a catalog of 174,397 journal articles that were published from 1993 through 2019, written in English, and tagged with the MeSH term <u>“computational biology”</u>, which is <u>equivalent</u> to “bioinformatics” and includes categories such as genomics and systems biology (via PubMed’s term explosion to include subterms). Excluding 163 articles with no author information and years with less than 200 articles/year, we analyzed 173,735 articles from 1998–2019.

### Countries of Afiliations

Publications often provide affiliation lists for authors, which generally associate authors with research organizations and their corresponding physical addresses. We implemented affiliation extraction in the pubmedpy Python package for both PubMed and PMC XML records. These methods extract a sequence of textual affiliations for each author.

We relied on two Python utilities to extract countries from text: <u>geotext</u> and <u>geopy.geocoders.Nominatim</u>. The first, geotext, used regular expressions to find mentions of places from the <u>GeoNames database</u>. To avoid mislabeling, we only mapped the affiliation to a country if geotext identified 2 or more mentions of that country. For example, in the affiliation string “Laboratory of Computational and Quantitative Biology, Paris, France”, geotext detected 2 mentions of places in France: Paris, France. In this case, we assign France to this affiliation.

This country extraction method accommodates multiple countries. Although ideally each affiliation record would refer to one and only one research organization, sometimes journals deposit multiple affiliations in a single structured affiliation. In these cases, we assigned multiple countries to the article. For more details on this approach, please consult the accompanying <u>notebook</u> and <u>label dataset</u>.

When geotext did not return results, we use the geopy approach, which returns a single country for an affiliation when successful. Its `geocoders.Nominatim` function converts names / addresses to geographic coordinates using the OpenStreetMap’s <u>Nomatim</u> service. With this method, we split a textual affiliation by punctuation into a list of strings and iterate backward through this list until we found a Nomatim search result. For the above affiliation, the search order would be “France”, “Paris”, and “Laboratory of Computational and Quantitative Biology”. Since Nomatim would return a match for the first term “France” (matched to France), the search would stop before getting to “Paris”, and “France” would be assigned to this affiliation.

Our ability to assign countries to authors was largely driven by the availability of affiliations. The country-assignment-rate for last authors from PubMed records was approximately 47%. This reflects the varying availability of affiliation metadata by journal.

For ISCB honorees, during the curation process, if an honoree was listed with their affiliation at the time, we recorded this affiliation for analysis. For ISCB Fellows, we used the affiliation listed on the ISCB page. Because we could not find affiliations for the 1997 and 1998 RECOMB keynote speakers’ listed for these years, they were left blank. If an author or speaker had more than one affiliation, each was inversely weighted by the number of affiliations that individual had.

### Estimation of Gender

We predicted the gender of honorees and authors using the https://genderize.io API, which was trained on over 100 million name-gender pairings collected from the web and is one of the three widely-used gender inference services [28]. We used author and honoree first names to retrieve predictions from genderize.io. The predictions represent the probability of an honoree or author being male or female. We used the estimated probabilities and did not convert to a hard group assignment. For example, a query to https://genderize.io on January 26, 2020 for “Casey” returns a probability of male of 0.74 and a probability of female of 0.26, which we would add for an author with this first name. Because of the limitations of considering gender as a binary trait and inferring it from first names, we only consider predictions in aggregate and not as individual values for specific scientists.

Of 412 ISCB honorees, genderize.io fails to provide gender predictions for one name. Of 173,735 last authors, 1,002 were missing a fore name in the raw paper metadata and 11,427 had fore names consisting of only initials. Specifically, the metadata for most papers before 2002 (2,566 out of 2,601 papers) only have initials for first and/or middle author names. Without gender predictions for these names, we consider only articles from 2002 on when comparing gender compositions between two groups. Of the remaining authors, genderize.io failed to predict gender for 9,845 of these fore names. We note that approximately 42% of these NA predictions are hyphenated names, which is likely because they are more unique and thus are more difficult to find predictions for. 82% of these names were predicted to be of Asian origin by last name (see the race/ethnicity prediction model below). This bias of NA predictions toward non-English names has been previously observed [29] and may have a minor influence on the final estimate of gender compositions.

### Estimation of Name Origin Groups

We developed a model to predict geographical origins of names. The existing Python package ethnicolr [30] produces reasonable predictions, but its international representation in the data curated from Wikipedia in 2009 [31] is still limited. For instance, 76% of the names in ethnicolr’s Wikipedia dataset are European in origin.

To address these limitations in ethnicolr, we built a similar classifier, a Long Short-term Memory (LSTM) neural network, to infer the region of origin from patterns in the sequences of letters in full names. We applied this model on an updated, approximately 4.5 times larger training dataset called Wiki2019 (described below). We tested multiple character sequence lengths and, based on this comparison, selected tri-characters for the primary results described in this work. We trained our prediction model on 80% of the Wiki2019 dataset and evaluated its performance using the remaining 20%. This model, which we term Wiki2019-LSTM, is available in the online file `LSTM.h5`.

To generate a training dataset for name origin prediction that reflects a modern naming landscape, we scraped the English Wikipedia’s category of <u>Living People</u>. This category, which contained approximately 930,000 pages at the time of processing in November 2019, is regularly curated and allowed us to avoid pages related to non-persons. For each Wikipedia page, we used two strategies to find a full birth name and location context for that person. First, we looked for nationality mention in the first sentence in the body of the text. In most English-language biographical Wikipedia pages, the first sentence usually begins with, for example, “John Edward Smith (born 1 January 1970) is an American novelist known for …” This structure comes from editor <u>guidance on biography articles</u> and is designed to capture:

> … the country of which the person is a citizen, national or permanent resident, or if the person is notable mainly for past events, the country where the person was a citizen, national or permanent resident when the person became notable.

Second, if this information is not available in the first sentence of the main text, we used information from the personal details sidebar; the information in this sidebar varied widely but often contained a full name and a place of birth.

We used regular expressions to parse out the person’s name from this structure and checked that the expression after “is a” matched a list of nationalities. We were able to define a name and nationality for 708,493 people by using the union of these strategies. This process produced country labels that were more fine-grained than the broader patterns that we sought to examine among honorees and authors. We initially grouped names by continent, but later decided to model our categorization after the hierarchical taxonomy used by <u>NamePrism</u> [32]. The NamePrism taxonomy was derived from name-country pairs by producing an embedding of names by Twitter contact patterns and then grouping countries using the similarity of names from those countries. The countries associated with each grouping are shown in Fig 5. NamePrism excluded the US, Canada and Australia because these countries have been populated by a mix of immigrant groups [32].

**Figure 5:**
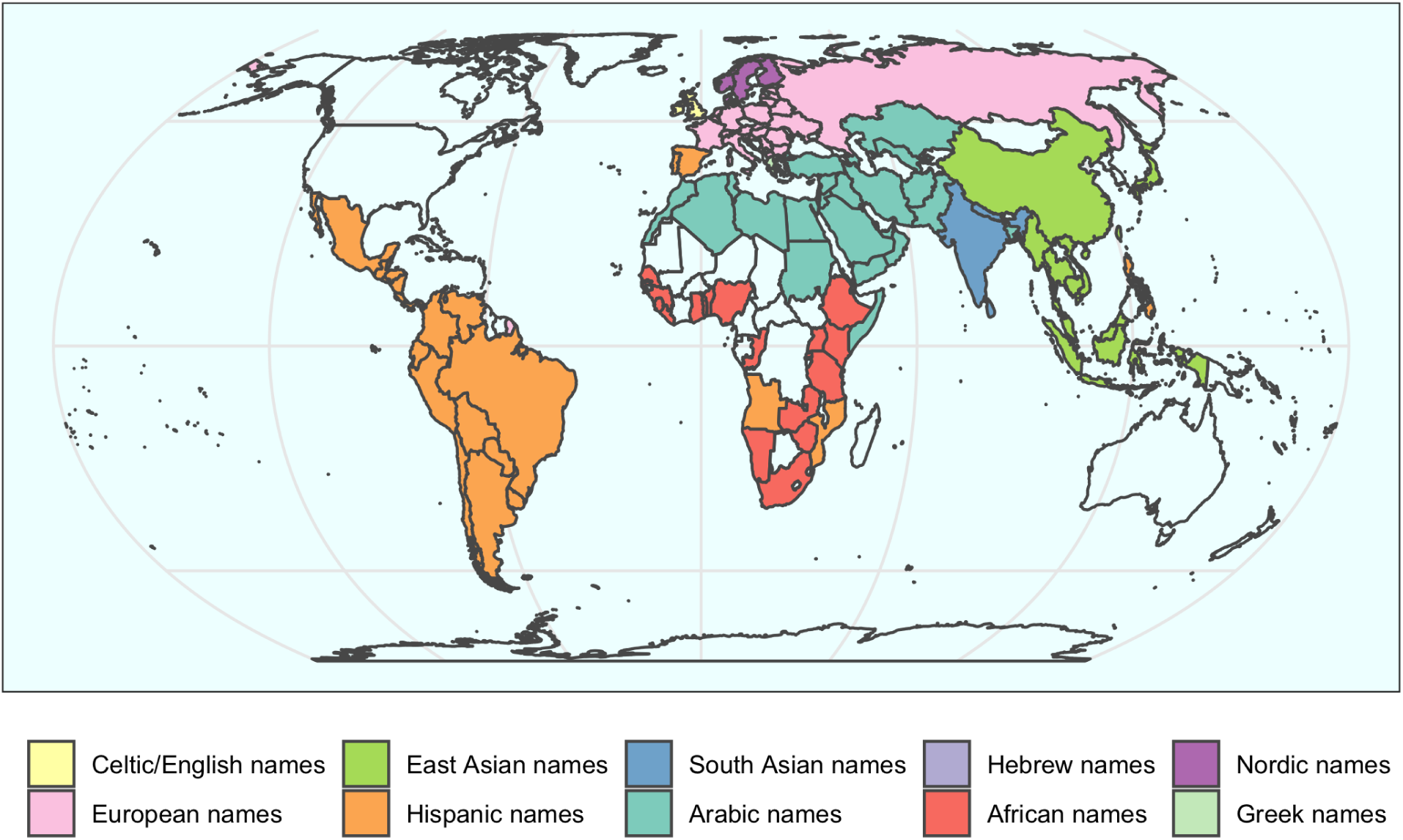
NamePrism groups countries by name similarity. We used this grouping but renamed the groups to focus on the linguistic patterns based on name etymology identified by NamePrism.

In an earlier version of this manuscript, we also used category names derived from NamePrism, but a reader <u>pointed out</u> the titles of the groupings were problematic; therefore, in this version, we renamed these groupings to reflect that the NamePrism approach primarily identifies groups based on linguistic patterns from name etymology rather than religious or racial similarities. We note that our mapping from nationality to name origins was not without error. For example, a scientist of Israeli nationality may not bear a Hebrew name. These mismatches were assessed via the heatmap of the model performance (Fig. 6C) and complemented by the affiliation analysis below. An alternative approach is to assign arbitrary names to these groups such as via letter coding (e.g., A, B, C, etc.), but we did not choose this strategy because ten arbitrary letters for ten groups can greatly reduce the paper’s readibility.

**Figure 6:**
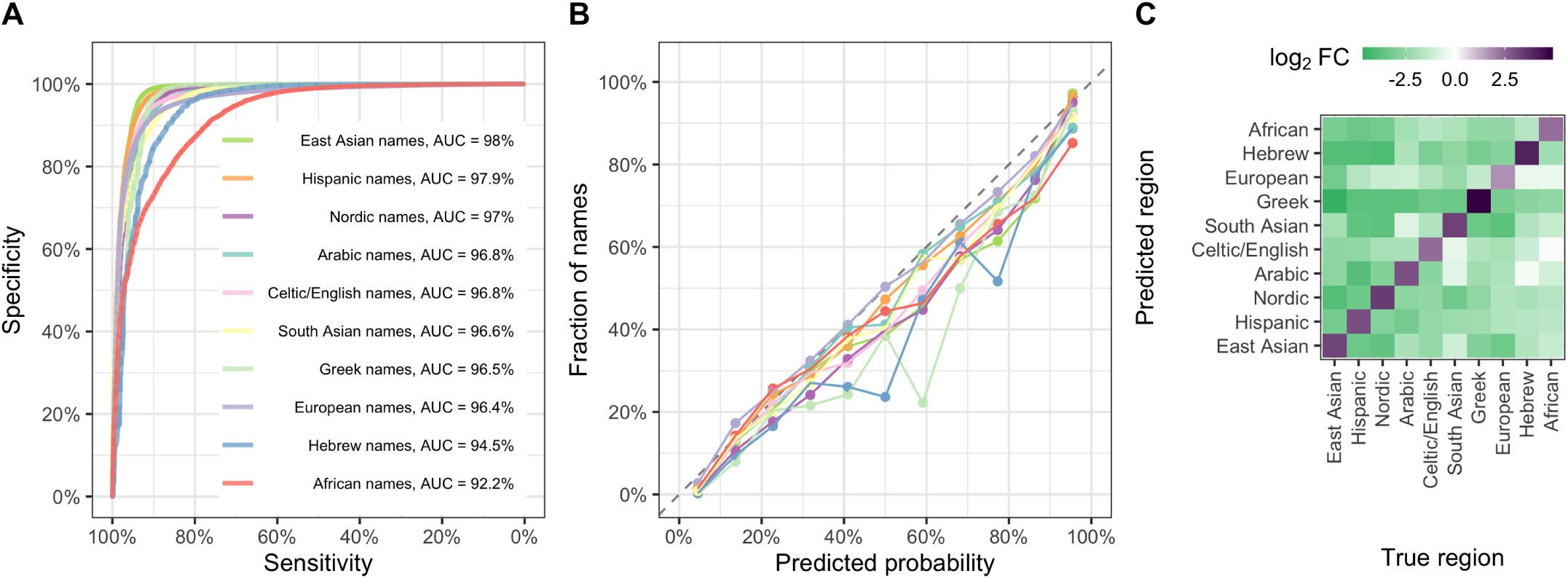
The Wiki2019-LSTM model performs well on the testing dataset. The area under the ROC curve is above 92% for each category, showing strong performance across origin categories (A). A calibration curve, computed with the caret R package, shows consistency between the predicted probabilities (midpoints of each fixed-width bin) and the observed fraction of names in each bin (B). Heatmap showing whether names from a given group (x-axis) received higher (purple) or lower (green) predictions for each group (y-axis) than would be expected by group prevalence alone (C). The values represent log_2_ fold change between the average predicted probability and the prevalence of the corresponding predicted group in the testing dataset (null). Scaling by group prevalence accounts for the imbalance of groups in the testing dataset. In all cases, the classifier predicts the true groups above the expected null probability (matrix diagonals are all purple). For off-diagonal cells, darker green indicates a lower mean prediction compared to the null. For example, the classifier does not often mistake East Asian names as Greek, but is more prone to mistaking South Asian names as Celtic/English.

### Predicting Name Origin Groups with LSTM Neural Networks and Wikipedia

Table 1 shows the size of the training set for each of the name origin groups as well as a few examples of PubMed author names that had at least 90% prediction probability in that group. We refer to this dataset as Wiki2019 (available online in <u>annotated_names.tsv</u>).

We next aimed to predict the name origin groups of honorees and authors. We constructed a training dataset with more than 700,000 name-nationality pairs by parsing the English-language Wikipedia. We trained a LSTM neural network on n-grams to predict name groups. We found similar performance across 1, 2, and 3-grams; however, because the classifier required fewer epochs to train with 3-grams, we used this length in the model that we term Wiki2019-LSTM. Our Wiki2019-LSTM returns, for each given name, a probability of that name originating from each of the specified 10 groups. We observed a multiclass area under the receiver operating characteristic curve (AUC) score of 95.9% for the classifier, indicating that the classifier can recapitulate name origins with high sensitivity and specificity. For each individual group, the high AUC (above 92%, Fig. 6A) suggests that our classifier was suffcient for use in a broad-scale examination of disparities. We also observed that the model was well calibrated (Fig. 6B). We also examined potential systematic errors between pairs of name origin groupings with a confusion heatmap and did not find off-diagonal enrichment for any pairing (Fig. 6C).

Applying Wiki2019-LSTM on the author and honoree datasets, we obtained name origin estimates for all honorees’ and authors’ name, except the 12,429 that did not have fore names (see breakdown in the Estimation of Gender section above). Once again, because the large majority of author fore names prior to 2002 were recorded with initials only, predictions were not possible, and we excluded 1998–2001 when comparing name origin compositions between two groups.

### Afiliation Analysis

For each country, we computed the expected number of honorees by multiplying the proportion of authors whose affiliations were in that country with the total number of honorees. We then performed an enrichment analysis to examine the difference in country affiliation proportions between ISCB honorees and field-specific last authors. We calculated each country’s enrichment by dividing the observed proportion of honorees by the expected proportion of honorees. The 95% confidence interval of the log_2_ enrichment was estimated using the Poisson model method [33].

### Estimation of Race and Ethnicity within the US

The underlying data used for race and ethnicity prediction are derived from the US Census, in which an individual’s race and ethnicity are based on their self-identification with one or more groups. Specifically, the race categories include White, Black or African American, American Indian or Alaska Native, Asian, Native Hawaiian or Other Pacific Islander, Other race, and Two or more races [34], and ethnicity categories include Hispanic/Latino or Not Hispanic/Latino [35]. We made race and ethnicity predictions of the surnames of US-affiliated authors and honorees using the R package wru, which implements methods described in Imai and Khanna [36]. wru uses similar race/ethnicity categories as in the Census but groups American Indian or Alaska Native and Native Hawaiian or Other Pacific Islander to form the Other category. In the case of names that were not observed in the census, wru’s *predict_race()* function outputs the average demographic distribution from the census, which may produce misleading results. To avoid this suboptimal imputation, we modified the function to return a status denoting that results were inconclusive (NA) instead. This prediction represents the probability of an honoree or author selecting a certain race or ethnicity on a census form if they lived within the US.

Of 239 US-affiliated ISCB honoree entries, wru fails to provide race/ethnicity predictions for 45 names. Of 26,771 US-affiliated last authors, 5,020 had a last name for which wru did not provide predictions. One limitation of wru and other methods that infer race, ethnicity, or nationality from last names is the potentially inaccurate prediction for scientists who changed their last name during marriage, a practice more common among women than men.

### Statistical analysis

We estimated the levels of representation by performing the following regression of the prediction probability on the group of scientists while controlling for year:

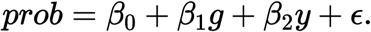

The dependent variable *prob* is the prediction probablity of a demographic variable (gender, race/ethnicity, and name origin) for names of scientists in *g* (honoree or author) during year *y. ϵ* ∼ *N*(0, *σ*^2^) accounts for random variation.

## Data and Resource Availability

This manuscript was written <u>openly on GitHub</u> using Manubot [37]. The Manubot HTML version is available under a Creative Commons Attribution (CC BY 4.0) License at https://greenelab.github.io/iscb-diversity-manuscript/. Our analysis of authors and ISCB-associated honorees is available under CC BY 4.0 at https://github.com/greenelab/iscb-diversity, with source code also distributed under a BSD 3-Clause License. Rendered Python and R notebooks from this repository are browsable at https://greenelab.github.io/iscb-diversity/. Our analysis of PubMed, PubMed Central, and author names relies on the Python pubmedpy package, developed as part of this project and available under a Blue Oak Model License 1.0 at https://github.com/dhimmel/pubmedpy and on <u>PyPI</u>. Our Wikipedia name dataset is dedicated to the public domain under CC0 License at https://github.com/greenelab/wiki-nationality-estimate, with source code to construct the dataset available under a BSD 3-Clause License.

